# Self–Terminating Bilayer Hydrogel Actuators via Enzyme–Programmed Mechanical Feedback

**DOI:** 10.64898/2026.03.10.710804

**Authors:** Fan Mo, Gali Bar–Shalom, Ehud Shimko Gozlan, Yuchen Liu, Alejandro Sosnik, Luai. R. Khoury

## Abstract

Autonomous soft materials that can actuate, perform a function, and then self–terminate without external intervention remain difficult to realize. Here, a bilayer hydrogel actuator fabricated by digital light processing-based 3D bioprinter is introduced that couples rapid thermoresponsive deformation with slower enzyme–programmed mechanical feedback to achieve self–regulated shape transformation and autonomous recovery. The system integrates a poly(N–isopropylacrylamide) actuation layer with a bovine serum albumin–poly(ethylene glycol) diacrylate enzyme–programmed layer loaded with trypsin. Above the lower critical solution temperature, deswelling of the actuation layer generates a strain mismatch across the bilayer and drives rapid closure. In parallel, proteolytic cleavage of albumin domains progressively softens the enzyme–programmed layer, reduces interlayer constraint, and acts as an intrinsic mechanical off–switch that relaxes curvature and restores the open state. This materials logic enables sustained enzyme release, time–dependent modulus loss, and autonomous shape recovery without staged external triggers. As a proof–of–concept, this platform is implemented as a gastrointestinal–retentive hydrogel gripper for localized intestinal enzyme delivery, where it exhibits thermally triggered gripping, millinewton–scale gripping force, autonomous reopening, and robust *ex vivo* retention on porcine small intestine under dynamic motion. These findings establish enzyme–programmed mechanical feedback as a general design strategy for self–regulated soft actuators and therapeutic materials with built–in functional lifetimes.

## Introduction

The gastrointestinal (GI) tract is essential for nutrient digestion and absorption, with the small intestine serving as the primary site of enzymatic breakdown and uptake [1]. Trypsin is a central pancreatic protease, secreted as trypsinogen and activated at the duodenal brush border by enterokinase (enteropeptidase) [2]. Beyond direct proteolysis, trypsin acts as a regulatory hub by activating multiple downstream digestive enzymes [3,4]. However, trypsin activity and/or concentration decrease with aging [5] and in medical conditions such as pancreatic insufficiency and GI infection [6], contributing to maldigestion and related disorders. Together, these limitations motivate alternative strategies that enable localized, sustained enzyme supplementation directly in the small intestine, the physiological site of action.

Oral administration remains the most popular route for small–molecule drugs owing to minimal invasiveness, ease of use, and patient compliance [7]. In the case of protein–and peptide–based therapeutics, it has become very appealing especially for local delivery in the different regions of the GI tract, while for systemic therapies it remains challenging due to the harsh degrading conditions of the stomach and the gut which reduce their bioavailability in the bloodstream [8]. From a broader perspective, oral delivery is uniquely challenging: drugs must dissolve appropriately, remain stable and functional while traversing diverse GI environments, and act at specific regions [9]. These barriers are compounded by poor long–term adherence–chronic medication adherence is often ∼50% in many settings [8,10,11]. In this framework, intensive research has been dedicated to designing a plethora of carriers for oral administration, especially for peptide/protein [12–14], though most of them cannot control regional release [15].

Ingestible devices with extended GI residence offer a compelling solution by combining non–invasive administration with prolonged, site–specific therapeutic action [16]. Recent GI–retentive designs span electronic platforms and soft actuator–based systems, typically relying on chemo– or mechano–adhesion for anchoring [16,17]. Soft actuators, inspired by biological motion, are particularly attractive because they are lightweight, mechanically compliant, and highly shape–adaptive–properties well matched to dynamic mucosal tissues [18]. Hydrogels, especially stimulus–responsive networks, are prominent actuator materials due to their very good biocompatibility and programmable swelling/deswelling, enabling shape changes in response to pH, temperature, electric fields, enzymes, and other physicochemical and biological cues [19–30].

Hydrogel–based GI–retentive systems such as thera–grippers and swellable polymers have demonstrated enhanced local drug delivery [31,32]. Thermoresponsive grippers can autonomously close above a transition temperature and sustain drug release for days [33]; variants further incorporate magnetic guidance or rigid support layers to improve orientating, grasping and retention [34,35]. More broadly, diverse microgrippers and geometries–based GI–retentive devices (e.g., stellate or coiled forms) have achieved impressive bending, wrapping, and expansion behaviors [36–42]. Yet a key limitation persists: most systems execute single–stage, one–way (irreversible) deformation to anchor tissue, with limited ability to autonomously reverse deformation and disengage after completing their function. As a result, prolonged, uncontrolled tissue intervention can increase the risk of mucosal abrasion and obstruction, restricting translation in confined, motile intestinal environments. Therefore, embedding intrinsic “negative feedback” into device function–so that actuation is initiated by physiology and then self–terminates–would enable patient–independent, safer retention and release [43].

Protein–polymer hybrid materials provide a powerful route to such self–regulated behavior. Protein incorporation can improve biocompatibility, adhesion, swelling, and biodegradability, while introducing enzymatically addressable domains [44]. Notably, protease–protein interactions can serve as an internal biochemical program that remodels network structure and mechanics over time [45].

Here, we report on an autonomous, stimulus–responsive bilayer hydrogel actuator for trypsin delivery in the small intestine, fabricated by multilayer digital light processing (DLP) 3D printing (Figure 1a(i)). The device comprises a thermoresponsive poly(N–isopropylacrylamide) (PNIPAM) actuation layer and an enzyme–programmed layer (BSA–PEGDA) loaded with trypsin (BSA–PEGDA(Trypsin)) (Figure 1a(i)). DLP printing enables high–resolution architectures with rapid, design–flexible fabrication. At physiological temperature (37°C), PNIPAM deswells (lower critical solution temperature, LCST ∼32 ° C), generating a programmed bending motion because PNIPAM undergoes a larger volumetric change than the BSA–PEGDA layer, creating a swelling mismatch across the bilayer (Figure 1a(ii)). This mismatch is translated into curvature by the mechanical constraint between the two layers: bending proceeds until the actuation–driven deformation is balanced by the elastic resistance of the bilayer, which depends on the stiffness and thickness of each layer and how they share load within the composite structure.

**Figure 1.**
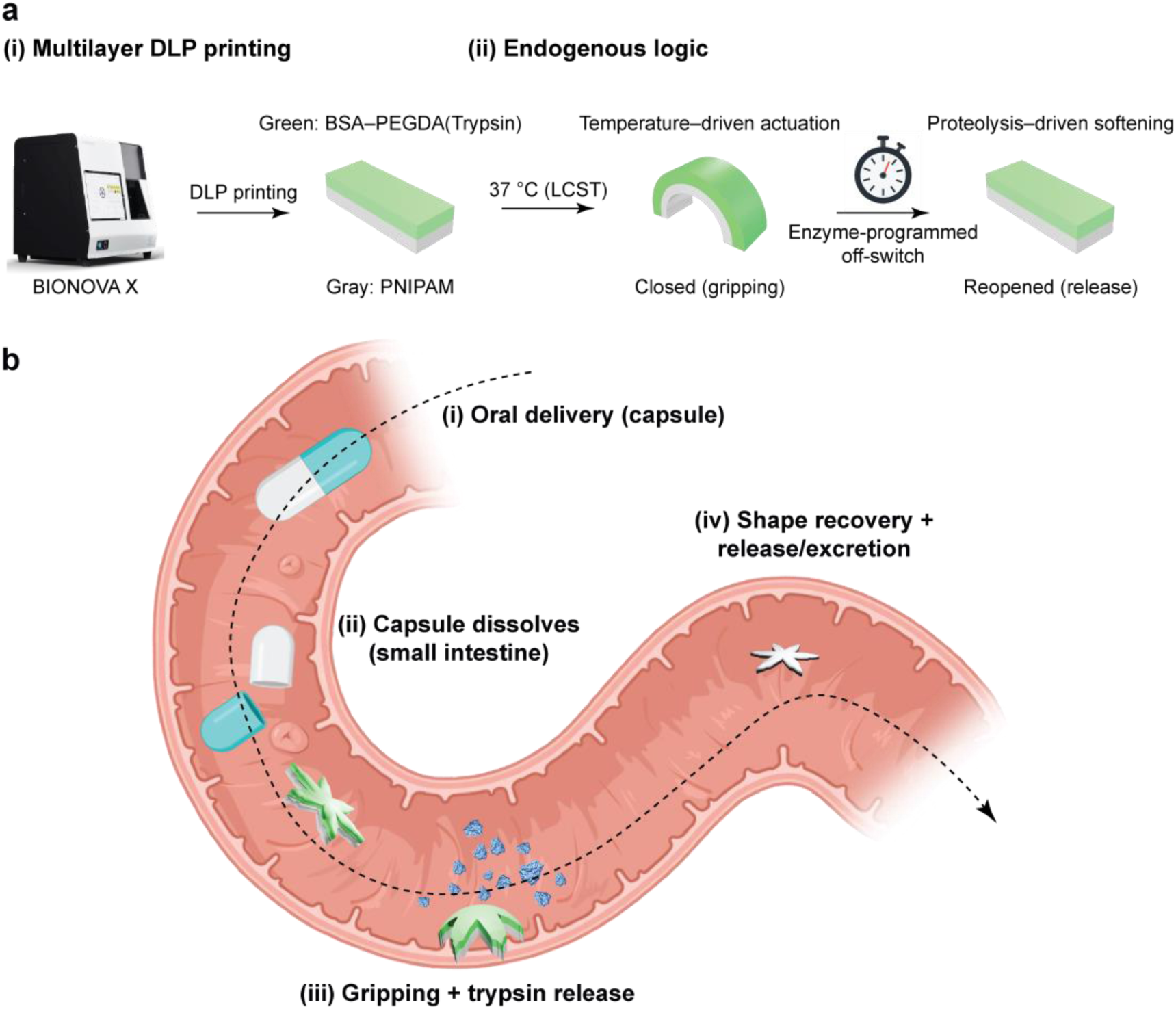
Concept and operating mechanism of autonomous thermoresponsive PNIPAM/BSA–PEGDA(Trypsin) bilayer hydrogel grippers for intestinal retention and enzyme delivery. **(a)** Multilayer DLP printing (BIONOVA X) of the bilayer (gray: PNIPAM; green: BSA–PEGDA(Trypsin)) and its endogenous actuation logic at intestinal conditions (37 °C, pH ∼8.0). Heating above the PNIPAM LCST drives deswelling-induced bending and gripper closure; concurrent proteolysis of BSA domains progressively softens the BSA–PEGDA layer, reducing interlayer constraint and triggering autonomous reopening (enzyme-programmed off-switch). **(b)** *In vivo*–mimicking sequence in the GI tract: (i). A pH-responsive capsule protects the gripper during gastric transit (pH ∼2.0) and (ii) dissolves in the small intestine (pH ∼8.0) to release the device. (iii). The gripper closes at 37 °C to anchor on mucosa and locally releases trypsin as the protein-containing layer undergoes proteolytic softening, (iv). continued degradation terminates actuation, restores the open configuration, and enables detachment and excretion.

In parallel, proteolytic digestion of BSA domains progressively softens the enzyme–programmed BSA–PEGDA(Trypsin) layer, enabling controlled trypsin release and–critically–providing an intrinsic mechanical “off–switch” that relaxes bending and promotes autonomous shape recovery and disengagement from the mucosal gripping site (Figure 1a(ii)). Mechanistically, proteolysis reduces the effective stiffness and load–bearing integrity of the BSA–PEGDA network (and can increase porosity and chain mobility), which weakens the bilayer’s ability to sustain curvature. As softening progresses, the stored elastic energy that maintains the closed configuration diminishes, curvature relaxes, and the interlayer constraint is effectively reduced–biasing the actuator back toward its low–curvature (open) state and enabling release.

Together, these coupled thermo–mechanical and enzyme–programmed mechanical changes establish a self–regulated GI–retentive bilayer hydrogel actuator that anchors under the stimulation of physiological temperature, delivers trypsin locally, and then autonomously releases via protease–induced softening behaviors (Figure 1b).

## Results

### DLP–printed PNIPAM/BSA–PEGDA bilayers: Actuation optimization and programmable geometries

PNIPAM (LCST ∼32°C) is well suited for thermally triggered actuation near physiological temperature [46–49]. PNIPAM layers were fabricated by DLP printing via photoinitiated polymerization of N–isopropylacrylamide (NIPAM) with N,N’ – methylenebis(acrylamide) (BIS) using lithium phenyl–2,4,6–trimethylbenzoylphosphinate (LAP) in Tris buffer (TRIS, pH ∼7.4), with tartrazine as a photoabsorber to control curing depth. To maximize thermally driven deformation for bilayer actuation, we quantified the weight–based relative shrinkage ratio (RSR (%) = (W_25_ – W_37_)/W_37_ × 100) of printed PNIPAM networks as a function of printing parameters and composition.

Printing temperature strongly affected thermoresponsive behavior: increasing the printing temperature from 25 to 35°C progressively reduced RSR of PNIPAM hydrogels (NIPAM–BIS = 2–0.05 M) (Figure 2a), consistent with printing near/above the LCST, which promotes chain association/collapse and yields a denser network with reduced swelling capacity [50]. However, it exhibited generally high RSR values (>300%) with no significant dependence on light exposure intensity between 50–100% (Figure 2b). Composition further modulated RSR (Figure 2c). The 2 – 0.05 M formulation showed the highest RSR (329 ± 57%), whereas increasing BIS (0.05 → 0.2 M) markedly decreased RSR, consistent with increased crosslink density restricting chain mobility and water uptake [51]. Lower NIPAM concentration (1 M) also reduced RSR compared with 2 M, likely due to reduced polymer content and a smaller thermally driven volume change. Based on these results, we selected NIPAM–BIS = 2–0.05 M, printed at 25°C and 75% exposure, as the optimal condition balancing printability and maximal thermoresponsive deformation for bilayer actuation.

**Figure 2.**
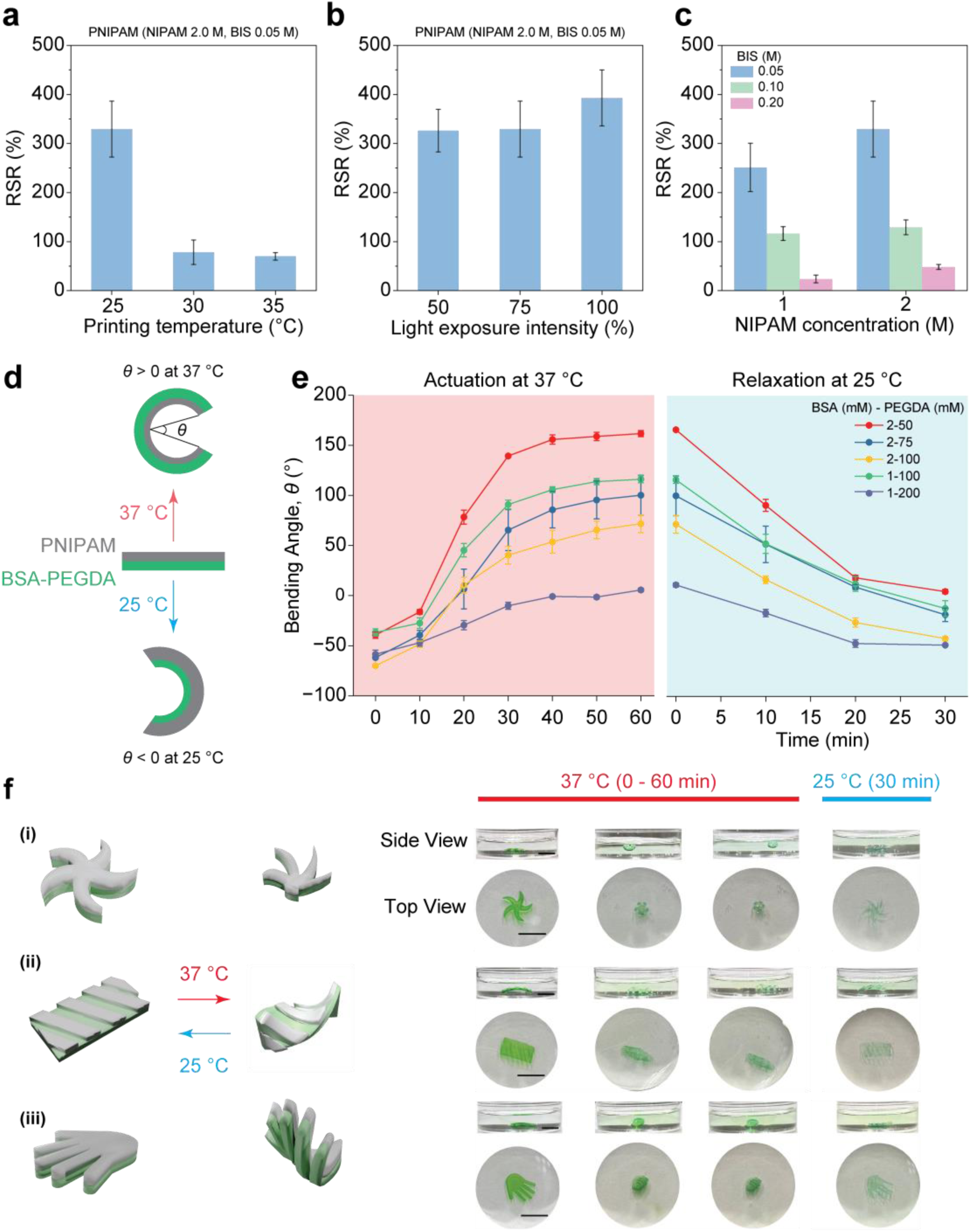
Printing parameter optimization and thermoresponsive actuation of DLP-printed PNIPAM/BSA–PEGDA bilayer actuators. **(a)** Effect of printing temperature (25–35 °C) on the relative shrinkage ratio (RSR) of PNIPAM (2 M NIPAM/0.05 M BIS) printed at 75% light intensity. **(b)** Effect of light exposure intensity (50–100%) on RSR for PNIPAM (2–0.05 M) printed at 25 °C. **(c)** Effect of formulation (NIPAM = 1 or 2 M; BIS = 0.05–0.2 M) on RSR (25 °C, 75% intensity). **(d)** Definition of bending angle (θ) for PNIPAM/BSA–PEGDA bilayers: PNIPAM swelling at 25 °C yields negative θ, while deswelling above the LCST of PNIPAM (37 °C) drives curvature reversal (positive θ). **(e)** Time-dependent bending at 37 °C (0.5 mm per layer) for bilayers with different BSA–PEGDA concentrations (BSA–PEGDA, mM–mM): 2–50, 2–75, 2–100, 1–100, and 1–200. Increasing BSA–PEGDA concentration reduces the final θ due to increased stiffness of the non-actuating layer. **(f)** Representative programmable shape morphing of PNIPAM/BSA–PEGDA (2.0/0.05 M; 2–50 mM–mM) in starfish, twisted, and hand geometries showing closure at 37 °C and recovery upon cooling to 25 °C. Scale bar: 6 mm. Data are mean ± SD (n = 3).

BSA–PEGDA hydrogels were prepared via aza–Michael coupling and subsequently DLP printed using LAP and tartrazine [24,52,53]. Bilayer actuators were then fabricated with PNIPAM as the thermoresponsive actuation layer and BSA–PEGDA as the enzyme–programmed structural layer for subsequent trypsin loading and mechanically regulated shape recovery. As illustrated in Figure 2d, at 25°C, PNIPAM is swollen, and the bilayer adopts a curved configuration; upon heating to 37°C, PNIPAM deswells and reverses curvature due to swelling mismatch.

We quantified thermal actuation by measuring the bending angle (θ) for PNIPAM (2–0.05 M)/BSA–PEGDA bilayers with varying BSA–PEGDA formulations (2–50, 2–75, 2–100, 1–100, 1–200 mM) and two per–layer thicknesses (0.25 and 0.5 mm) (Figure 2e; Figure S1-3). All actuators exhibited rapid bending during heating to 37°C and reversibility upon cooling to 25°C (∼30 min). Increasing BSA–PEGDA concentration decreased the final bending angle, consistent with a stiffer BSA–PEGDA layer resisting PNIPAM–driven deformation (cf. modulus trends with BSA–PEGDA composition) [30]. Thinner bilayers (0.25 mm per layer) produced larger bending amplitudes than thicker ones (0.5 mm per layer), but excessive curvature (>180°) was undesirable for stable thera–gripper retention (Figures S1–3). We therefore selected BSA–PEGDA (2–50 mM) with 0.5 mm per layer as the best compromise between bending amplitude and stable actuator geometry.

Finally, to demonstrate geometric versatility enabled by DLP printing and endogenous thermal triggering, we printed three representative bilayer architectures: a starfish gripper, a twisted sheet, and a hand–like actuator (Figure 2f). The starfish design transitioned from an expanded state to a closed configuration at 37°C, supporting a gripper concept for GI retention or sampling (Figure 2f(i)). The twisted bilayer, printed with an oblique PNIPAM orientation relative to the BSA–PEGDA layer, transformed into a tubular geometry upon heating, suggesting potential as a thermally triggered lumen–conforming structure (Figure 2f(ii)). The hand–like design closed into a fist at 37°C and reopened on cooling to room temperature (RT, below the LCST of PNIPAM), illustrating biomimetic grasping motions (Figure 2f(iii)). In a gradual heating/cooling protocol (25 → 37 → 25°C), all geometries completed shape morphing within ∼30 min and recovered within ∼30 min (Figure 2f). When the starfish actuator was placed directly into a preheated bath (37°C), closure accelerated to ∼5 min (Video S1), highlighting the potential of rapid thermal responsiveness under physiological temperature.

### Enzyme–programmed network softening and trypsin release from BSA–PEGDA hydrogels

To exploit protease–protein interactions as an internal “timer” for release and mechanical regulation, trypsin was physically encapsulated within the enzyme–programmed BSA–PEGDA network by mixing BSA–PEGDA with trypsin (25.2 μM) and LAP (4:1:1; v/v/v), yielding a physiologically relevant trypsin content (0.01% w/v, ∼4.2 μM) [54], followed by addition of tartrazine (1% w/v; 1:60, v/v) prior to DLP printing. Under stimulated intestinal conditions (TRIS, pH ∼ 8.0, 37°C), encapsulated trypsin progressively cleaved BSA domains, driving network softening/collapse while enabling enzyme release (Figure 3a).

**Figure 3.**
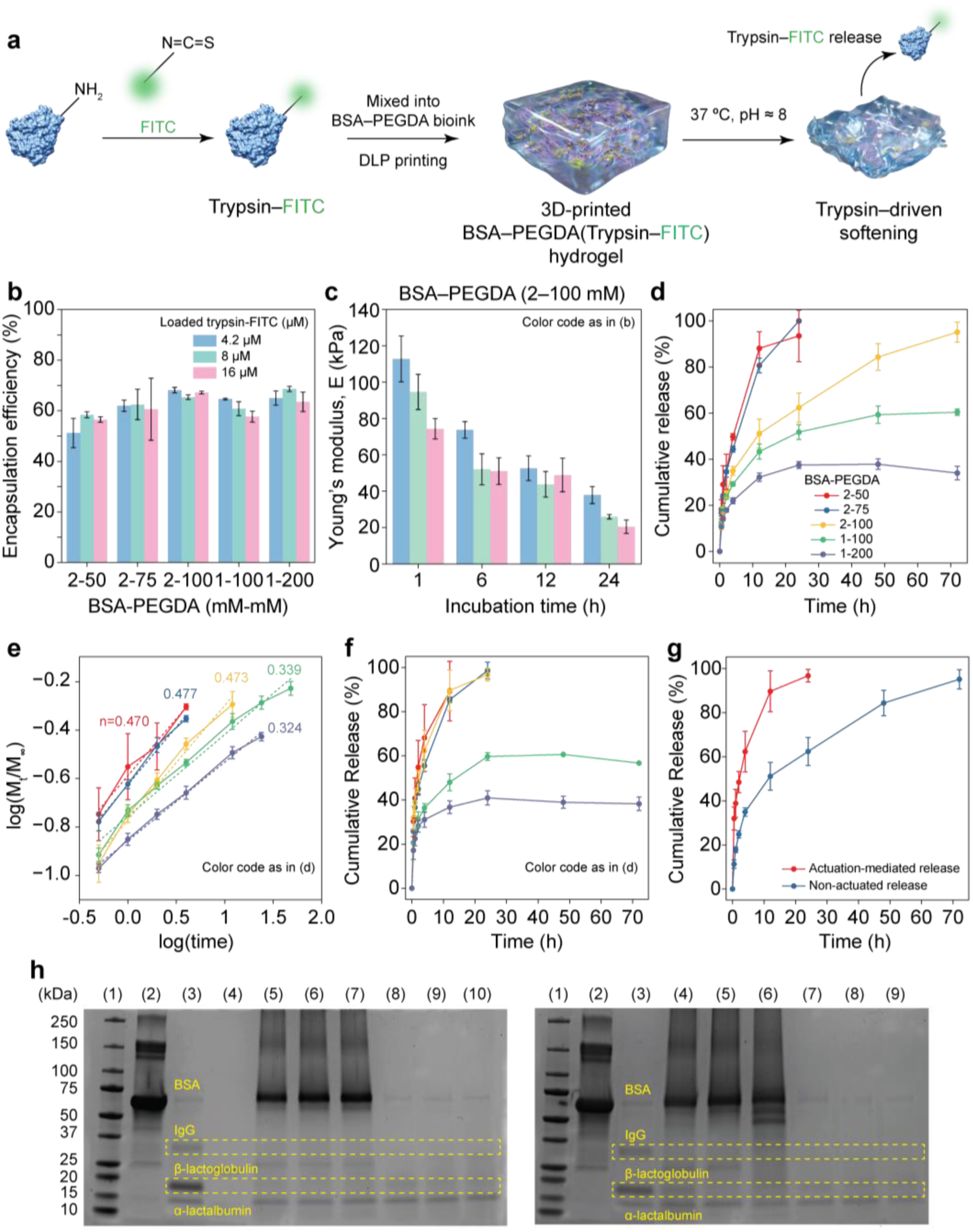
Encapsulation, release, and functional activity of trypsin–FITC from BSA–PEGDA hydrogels. **(a)** Schematic of FITC labeling of trypsin, encapsulation during DLP printing to form BSA–PEGDA(Trypsin–FITC) hydrogels, and proteolysis-driven network softening that promotes sustained release under intestinal-mimicking physiological conditions (37 °C, pH ∼8.0). **(b)** Encapsulation efficiency (EE) of trypsin–FITC (loaded: 4.2, 8, and 16 µM) in BSA–PEGDA formulations(BSA/PEGDA, mM–mM): 2–50, 2–75, 2–100, 1–100, and 1–200. **(c)** Young’s modulus of BSA– PEGDA (2–100) hydrogels loaded with trypsin (4.2, 8, or 16 µM) after incubation for 1–24 h (modulus calculated from 0–10% strain), showing time- and loading-dependent softening. **(d)** Cumulative trypsin–FITC release from cylindrical BSA–PEGDA hydrogels at 37 °C and pH ∼8.0 (non-actuated) and **(e)** corresponding KP fits (release exponent *n* indicated). **(f)** Cumulative release from PNIPAM/BSA–PEGDA bilayers during thermal actuation and **(g)** comparison of actuated vs non-actuated release for BSA–PEGDA (2–100), highlighting actuation-enhanced transport. **(h,i)** SDS–PAGE of whey protein digestion by trypsin released from BSA–PEGDA(Trypsin) hydrogels: **(h)** time dependence (48–96 h) for 2–50 and 1–200 (8 µM trypsin) ((1). Ladder; (2). BSA; (3). Whey protein; (4). Trypsin; (5). BSA-PEGDA (2-50) in 48 h; (6). BSA-PEGDA (2-50) in 72 h; (7). BSA-PEGDA (2-50) in 96 h; (8). BSA-PEGDA (1-200) in 48 h; (9). BSA-PEGDA (1-200) in 72 h; (10). BSA-PEGDA (1-200) in 96 h) and **(i)** effect of trypsin loading (4.2–16 µM; 72 h) for 2–50 and 1–200 ((1). Ladder; (2). BSA; (3). Whey protein; (4). BSA-PEGDA (2-50) with encapsulated trypsin (4.2 μM); (5). BSA-PEGDA (2-50) with encapsulated trypsin (8 μM); (6). BSA-PEGDA (2-50) with encapsulated trypsin (16 μM); (7). BSA-PEGDA (1-200) with encapsulated trypsin (4.2 μM); (8). BSA-PEGDA (1-200) with encapsulated trypsin (8 μM); (9). BSA-PEGDA (1-200) with encapsulated trypsin (16 μM)). Disappearance of β-lactoglobulin and IgG bands indicates retained proteolytic activity. Data are mean ± SD (n = 3).

To quantify release, trypsin was fluorescently labeled with fluorescein isothiocyanate (FITC) to generate trypsin–FITC conjugates, which were then encapsulated to obtain BSA–PEGDA(Trypsin–FITC) hydrogels. Encapsulation efficiency (EE%) across formulations and loading levels (4.2, 8, 16 μM) was typically 55–75%, consistent with partial loss during post–printing washing (Figure 3b). Among the tested networks, 2–100 mM BSA–PEGDA showed the most uniform and highest retention across all loading concentrations relative to 2–50 and 2–75 mM, consistent with a denser network limiting washout. Additionally, EE% was higher for 2–100 mM than 1–100 mM, supporting the role of BSA as a reinforcing, covalently integrated component that enhances network integrity and enzyme retention (Figure 3b).

We next examined trypsin–programmed mechanical softening using 2–100 mM BSA–PEGDA(Trypsin) hydrogels loaded with 4.2, 8, or 16 μM trypsin and incubated at pH ∼8.0 and 37°C for 1–24 h (Figure 3c; Figure S4a–c). In all groups, Young’s modulus decreased monotonically with time, confirming progressive digestion of BSA crosslinking domains and loss of load–bearing structure. Higher trypsin loading produces faster and deeper softening. Notably, the 8 and 16 μM groups exhibited a transient plateau between 6–12 h, followed by renewed softening by 24 h, suggesting that local confinement within the dense network initially limits access to remaining cleavable domains, with additional degradation occurring as the network relaxes and fragmentation progresses. By 24 h, the 16 μM group approached ∼20 kPa, comparable to PEGDA–only networks [30], indicating near–complete loss of BSA–derived crosslinking contributions.

We then characterized trypsin–FITC release from cylindrical BSA–PEGDA hydrogels under simulated intestinal conditions (pH ∼8.0, 37°C) (Figure 3d; Figure S5). Lower–density networks (2–50 and 2–75 mM) released rapidly, reaching ∼100% within 24 h, whereas 2–100 mM exhibited a sustained profile extending to ∼72 h, consistent with increased compactness restricting transport. Interestingly, 1–100 mM showed slower release kinetics and lower cumulative release than 2–100 mM, despite its lower network density. This behavior is consistent with the BSA content acting not only as a structural element but also as the enzymatically cleavable program: higher BSA content (2–100 mM) provides more cleavage sites, promoting greater fragmentation and facilitating release, whereas 1–100 mM provides fewer cleavable domains and thus less degradation–assisted release. The 1–200 mM group showed the lowest release rate and incomplete cumulative release, consistent with a highly crosslinked network. Incomplete release for 1–100 and 1–200 mM may also reflect trypsin autolysis and/or partial fluorophore photobleaching during long incubations.

Release kinetics were fitted using the Korsmeyer–Peppas (KP) model (Figure 3e). For 2–50, 2–75, and 2–100 mM, the release exponent n fell between 0.45 and 1.0, indicating anomalous transport [55], consistent with combined diffusion and matrix relaxation/erosion driven by progressive enzymatic BSA digestion. To decouple enzymatic effects, release was also measured at 4°C (reduced trypsin activity) for 2–50, 2–75, and 2–100 mM (Figure S6). Under these conditions, release slowed down markedly: 2–50 mM approached completion only after ∼96 h, while 2–75 and 2–100 mM plateaued at ∼40% and ∼20%, respectively. KP fits (Figure S7) showed n decreased to 0.38 (2–75) and 0.319 (2–100), indicating a shift toward diffusion–driven (Fickian) release when enzyme–mediated network relaxation is suppressed.

To reflect device–relevant conditions, we further quantified release under bilayer actuation (Figure 3f). For 2–50 and 2–75 mM, actuation had little effect, consistent with their already rapid diffusion–driven release. In contrast, 2–100 mM exhibited a pronounced acceleration, reaching ∼100% release within 24 h under actuation (Figure 3g), with a smaller enhancement for 1–100 mM. This suggests a synergy between (i) actuation–induced mechanical stress that promotes transport (e.g., network stretching/extrusion) and (ii) the higher density of BSA cleavage sites in 2–100 mM that generates additional fragmentation pathways, together amplifying release in the actuated state (Figure S8).

Finally, we verified functional activity of released trypsin using digestion of whey proteins, analyzed by sodium dodecyl sulfate–polyacrylamide gel electrophoresis (SDS–PAGE) (Figure 3h). Whey bands were consistent with major components (β–lactoglobulin, α–lactalbumin, IgG fragments, and BSA) [56] (Figure 3h, lane 3). Upon incubation with BSA–PEGDA(Trypsin) hydrogels, β–lactoglobulin and IgG bands progressively diminished over 48–96 h, consistent with literature [57,58], whereas α–lactalbumin remained comparatively resistant, consistent with limited accessibility of trypsin cleavage sites in its compact fold and prior selective hydrolysis strategies [59–61]. The appearance/intensification of BSA bands in some conditions is consistent with trypsin–driven hydrogel degradation and diffusion of BSA–derived fragments into solution; in contrast, 1–200 mM remained structurally stable, consistent with minimal BSA band increase (Figure 3h). Varying encapsulated trypsin concentration (4.2, 8, 16 μM) increased digestion extent, most clearly reflected by reduced β–lactoglobulin intensity and increased low–M_w_ fragmentation in BSA–containing conditions (Figure 3i). Across BSA–PEGDA formulations, whey digestion remained effective (Figure S9), indicating that despite formulation–dependent release kinetics, the released trypsin amount was sufficient for proteolysis, with additional bands at ∼20–25 kDa consistent with peptide fragments produced during degradation of both hydrogel–derived and solution–phase proteins.

### Trypsin–programmed actuation termination and autonomous shape recovery

Hydrogel soft robotics offers biocompatible platforms capable of programmable deformation and adaptive responses [62]. Building on the trypsin–programmed softening and release described above, we next integrated this mechanism into bilayer actuators to enable autonomous actuation termination and shape recovery. PNIPAM/BSA–PEGDA(Trypsin) bilayers initially exhibited strong thermally driven bending at 37°C, followed by gradual reopening during long–term incubation under simulated intestinal conditions (TRIS, pH ∼8.0, 37°C). This recovery arises from progressive trypsin–mediated degradation/softening of the enzyme–programmed BSA–PEGDA layer, which diminishes the interlayer constraint required to sustain curvature and relaxes the actuator toward its low–curvature (open) configuration. From a bilayer–mechanics perspective, the closed state is maintained by a balance between the swelling–mismatch–generated curvature and the composite bending stiffness of the laminated structure. Proteolysis reduces the effective modulus and load–bearing continuity of the BSA–PEGDA layer (via cleavage of BSA domains and network fragmentation), thereby lowering the bilayer bending stiffness and weakening stress transfer across the interface. As a result, the bilayer can no longer sustain the elastic bending moment required to hold the closed configuration, and curvature progressively relaxes toward the open state.

To determine how the enzyme–programmed layer composition governs recovery kinetics, we fabricated PNIPAM/BSA–PEGDA(Trypsin) actuators with BSA–PEGDA formulations of 2–50, 2–75, 1–100, and 2–100 mM (each loaded with 4.2 μM trypsin) and monitored bending angle over one month (Figure 4a). The 2–50 mM group exhibited the earliest and fastest loss of bending capacity, with a pronounced decrease beginning on day 3 and reaching 28 ± 6° by day 30. This accelerated opening is consistent with the combined effects of enzymatic cleavage and the lower structural stability of the weaker crosslinked network. Notably, prior work reported that 2–50 mM BSA–PEGDA hydrogels retain structure for ∼7 days in TRIS and fully degrade after ∼45 days [30], indicating that encapsulated trypsin substantially accelerates network breakdown and mechanically programmed recovery. The 2–75 mM group showed a delayed but clear decline, initiating around day 7 and decreasing more gradually to 48 ±10° by day 30, consistent with increased resistance to digestion in a more compact network. In contrast, 1–100 and 2–100 mM actuators maintained nearly constant bending angles over 30 days, indicating that these stiffer, more densely functionalized networks were not sufficiently degraded by 4.2 μM encapsulated trypsin on this timescale (Figure 4a; Figure S10). Mechanistically, increasing BSA–PEGDA concentration increases the stiffness and structural redundancy of the enzyme–programmed layer, raising the mechanical barrier for curvature relaxation; therefore, more extensive protease cleavage (or longer time and/or additional protease) is required before the bilayer loses enough constraint to reopen.

**Figure 4.**
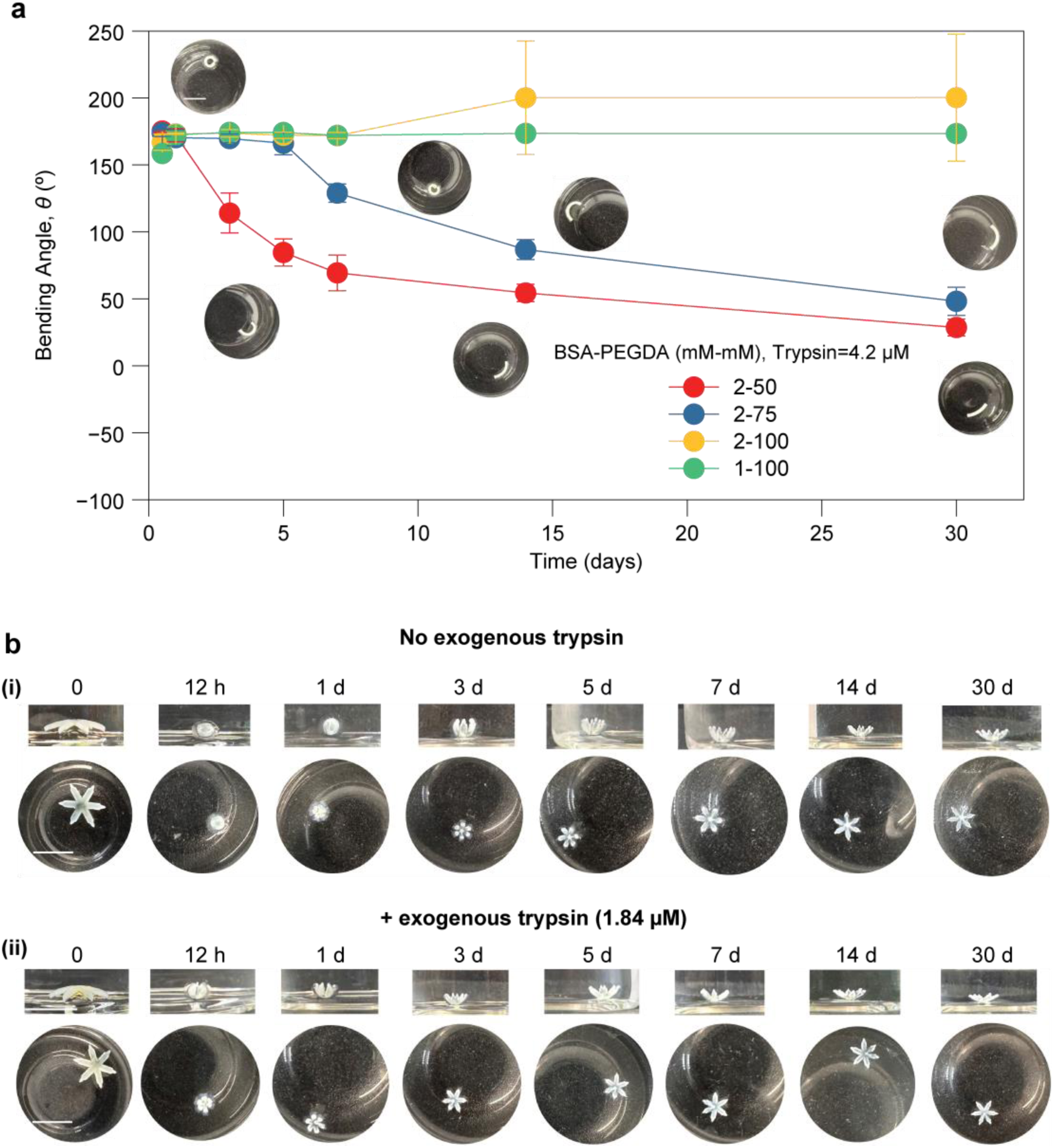
Autonomous shape recovery of PNIPAM/BSA–PEGDA(Trypsin) bilayer actuators via proteolysis-programmed softening. **(a)** Time-dependent bending angle (θ) during incubation at 37°C (pH ∼8.0) for bilayers with BSA–PEGDA(Trypsin) layers of different formulations (BSA/PEGDA, mM–mM): 2–50, 2–75, 1–100, and 2–100. Softer networks (2–50 and 2–75) exhibit progressive loss of curvature (opening), while stiffer formulations (1–100 and 2–100) maintain near-constant θ over the same period. **(b)** Representative time-lapse images of gripper opening in (i) TRIS (pH ∼8.0, 37°C) and (ii) a trypsin-insufficiency–mimicking condition (exogenous trypsin added at reduced concentration; 1.84 μM), illustrating accelerated reopening/detachment under proteolytic conditions. Scale bars: 6 mm. Data are mean ± SD (n = 3).

We therefore increased encapsulated trypsin in the 2–100 mM formulation to 8 and 16 μM (Figure S11a). Despite the higher loading, bending angles remained largely unchanged over the first 30 days, with only gradual opening detected after ∼60 days, suggesting that enzyme access and/or mobility within the dense network limits degradation–driven recovery on shorter timescales. In this regime, the rate–limiting step is likely not catalytic activity but the ability of trypsin to access cleavable BSA domains within a compact network and to diffuse as the matrix evolves, delaying the point at which the bilayer’s effective stiffness drops below the threshold needed to maintain curvature. During the early stage of enzymatic digestion, the polymer network became progressively loosened due to active enzyme cleavage. Although the enzymatic activity gradually diminished over time, the cumulative structural degradation continued to increase with the incubation periods, resulting in a progressive weakening of the hydrogel network after ∼60 days. To emulate a more protease–rich environment, actuators of the same formulation were incubated in an external simulated physiological trypsin solution (4.2 μM) (Figure S11b). Compared with incubation without external trypsin, these actuators exhibited markedly enhanced opening, and both conditions converged to ∼90° by ∼2 months, confirming that exogenous trypsin accelerates network degradation and promotes shape recovery.

Finally, we demonstrated practical self–release using a six–arm snowflake gripper fabricated from PNIPAM/BSA–PEGDA(Trypsin) (2–0.05 M with 2–50 mM BSA–PEGDA, loaded with 4.2 μM trypsin), selected for its robust recovery behavior in the bilayer screening (Figure S12). In TRIS (pH ∼8.0, 37°C), grippers began opening on day 3 and achieved complete release by day 14 (Figure 4b(i)). To better mimic reduced trypsin levels reported in insufficiency [63], grippers were incubated in a below–physiological trypsin solution (trypsin insufficient condition, ∼1.84 μM). Under these conditions, opening initiated on day 1 and complete release occurred by day 7 (Figure 4b(ii)), demonstrating that even modest exogenous protease levels can substantially accelerate programmed self–release. Mechanistically, external trypsin increases the effective proteolytic flux at the hydrogel surface and within accessible pores, accelerating cleavage–driven softening and thereby shortening the time required to reduce bilayer constraint below the curvature–sustaining threshold.

To further confirm the applicability and feasibility of the autonomous recovery behavior of hydrogel grippers, we tested in a more realistic scene. The as–developed PNIPAM/BSA–PEGDA(Trypsin) bilayer grippers were immersed in a high–concentration trypsin solution (∼1% w/v), with TRIS (37°C, pH ∼8.0) serving as the control group. In both cases, the grippers were initially anchored onto a custom–designed and 3D–printed platform that contained a ball as a grasp site. After 24 h of incubation, grippers exposed to trypsin solution gradually reopened and ultimately detached from the platform, whereas no comparable structural recovery was observed in the control group, demonstrating the significance of enzyme–triggered macroscopic reconfiguration on bilayer hydrogel grippers (Video S2). Therefore, the developed hydrogel gripper is not restricted to trypsin delivery but represents a versatile platform for GI therapy, capable of accommodating diverse therapeutic cargos (e.g., doxorubicin) while exploiting endogenous enzymatic activity as an intrinsic trigger for controlled release.

### Gripping force calibration, cell viability characterization, and *ex vivo* retention on porcine small intestine

To quantify gripping performance and establish a baseline for future *in vivo* studies, we developed a simple force–calibration assay based on magnet–assisted lifting (Figure S13). Using PNIPAM/BSA–PEGDA (2–0.05 M and 2–50 mM) bilayer actuators, the bending angle progressively decreased as the applied load increased from 0 to 20 magnets (step = 4 magnets) (Figure S13). At 16 magnets, the actuator opened to 92±3° (∼90°), indicating substantial loss of gripping capacity and corresponding to a measured force of 1.03 mN. We therefore define the characteristic gripping force of our bilayer actuator platform as ∼1 mN under these conditions (Figure 5a; Figure S13).

**Figure 5.**
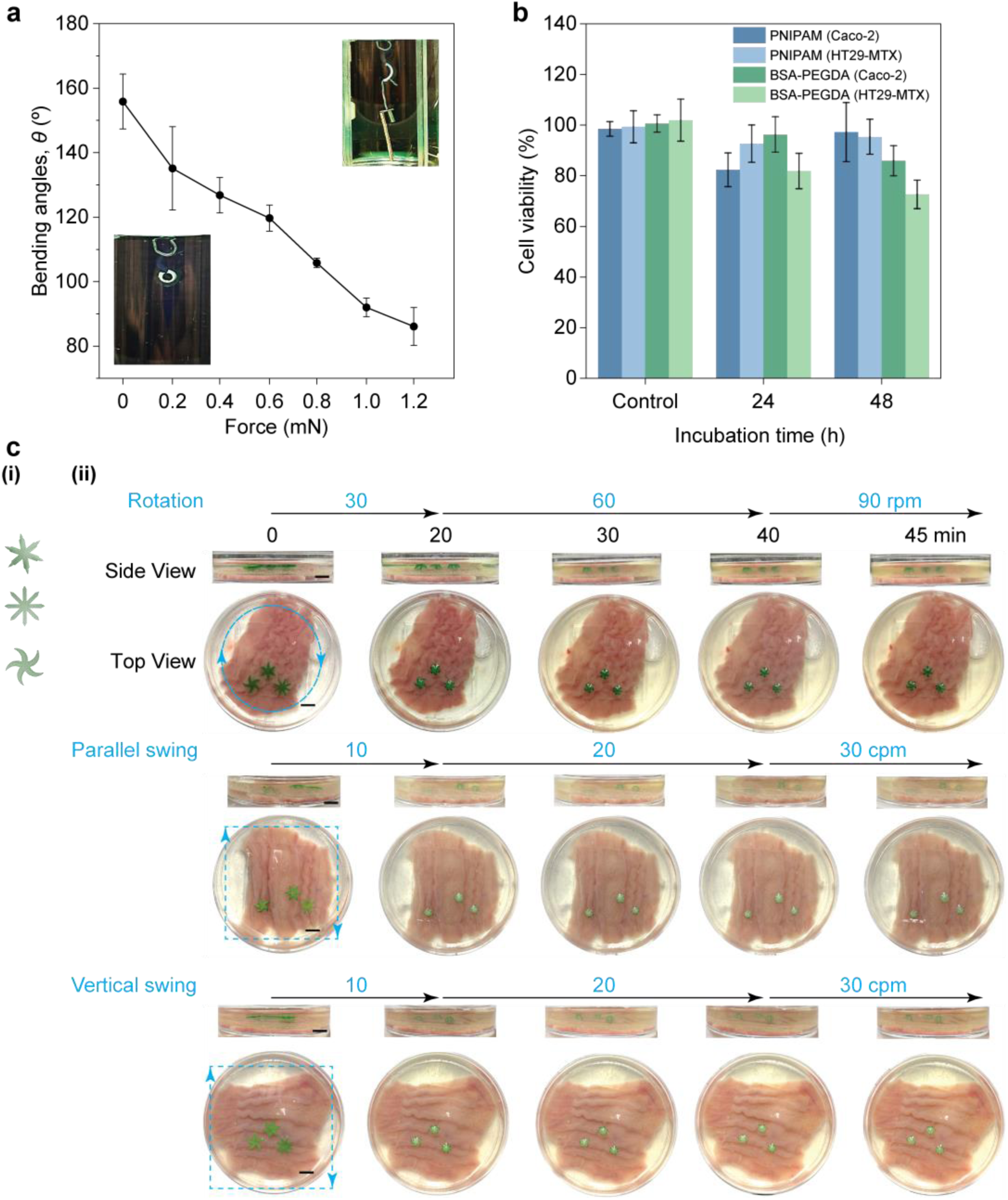
Gripping force, cytocompatibility, and *ex vivo* grasping of PNIPAM/BSA–PEGDA bilayer grippers. **(a)** Gripping force calibration for PNIPAM/BSA–PEGDA (2 – 0.05 M/2–50 mM–mM) actuators using incremental magnetic loading (0–20 magnets; +4 per step); increasing load reduces the bending angle toward ∼90°, defining the characteristic gripping force. **(b)** Cytocompatibility of PNIPAM (2–0.05 M) and BSA–PEGDA (2–50 mM–mM) hydrogels with Caco-2 and HT29-MTX cells after 24 and 48 h (viability >70%). **(c)** Representative gripper geometries (starfish, 6-arm edged snowflake, and 8-arm snowflake) and corresponding *ex vivo* grasping on porcine small intestine under dynamic motion: rotation (30 → 60 rpm; 90 rpm challenge) and swing (10 → 20 cpm; 30 cpm challenge) in directions parallel or perpendicular to plicae circulares. Scale bar: 6 mm. Data are mean ± SD (n = 3).

Biocompatibility is a critical consideration in the development of medical devices, particularly for GI retentive systems in which prolonged exposure is envisioned. The GI tract is covered by a mucous layer that protects the underlying epithelium from physical, chemical, biological and mechanical injury [64,65]. To evaluate cytocompatibility, cell viability was assessed for each hydrogel (PNIPAM and BSA–PEGDA hydrogels) (Figure 5b). To better mimic the epithelial microenvironment of the small intestine, cytocompatibility was evaluated using mucus–secreting human intestinal epithelial HT29–MTX cells and colorectal adenocarcinoma–derived Caco–2 cells, both well–accepted intestinal models *in vitro* [66]. Cell viability was estimated via the 3–(4,5–dimethylthiazol–2–yl)–2,5–diphenyltetrazolium bromide (MTT) assay. As the tissue–contacting layer of the bilayer gripper, PNIPAM hydrogels exhibited high cell viability after 24 h (∼82% for Caco–2 and ∼92% for HT29–MTX) and 48 h (∼97% and ∼95%, respectively). Similarly, BSA–PEGDA hydrogels maintained excellent viability (>80%) in Caco–2 cells, while slightly reduced viability was observed in HT29–MTX cells, particularly at 48 h (∼70%). Nevertheless, all the cell viability values remained above the generally accepted range for cytocompatible biomaterials *in vitro* [67]. Collectively, these results confirm the compatibility of both hydrogel components for bilayer actuator fabrication. We next evaluated practical gripping on fresh porcine small intestine as an *ex vivo* model due to its anatomical and physiological similarity to the human intestine [68]. For this, fresh small intestines were obtained from healthy adult pigs at a White Millya slaughterhouse (Israel) within 2 h postmortem according to animal protection regulations, and without fasting them prior to sample collection. Because gripper geometry governs how forces are distributed at the tissue–device interface and can strongly affect anchoring efficiency, we compared three designs: a starfish, an 8–arm snowflake, and a 6–arm edged snowflake (Figure 5c(i)). The starfish geometry was selected to promote rotational engagement during actuation, while the 8–arm snowflake tested whether increasing arm number within design constraints improves tissue grasping. Inspired by reports that sharper distal features can possibly enhance local anchoring [69], we additionally designed the 6–arm edged snowflake to increase localized contact and tissue engagement.

To better approximate intestinal dynamics, we tested retention under three motion modes representing distinct flow/loading directions relative to the porcine small–intestine surface morphology: rotation, plicae circulares–parallel swing, and plicae circulares–vertical swing (Figure 5c(ii)) [70]. Porcine intestine segments were fixed in a custom resin 3D–printed holder (Figure S14), grippers were placed onto the mucosal surface in pre–heated solution, and controlled rotation/swing was applied. To allow stable initial anchoring, tests began with gentle motion (30 revolutions per minute (rpm); 10 cycles per min (cpm) swing) for 20 min, followed by a sustained testing stage (60 rpm; 20 cpm) for an additional 20 min, and finally an extreme challenge step (90 rpm; 30 cpm) for 1 min.

Across geometries and motion modes, grippers gradually established contact and grasp during the initial low–stress phase, then maintained stable attachment throughout the 20–40 min testing interval under both rotation and swing conditions (Figure 5c(ii)). Even during the extreme challenge step, gripper position and configuration remained largely unchanged (Figure 5c(ii)). We attribute this performance to the combination of thermally triggered closure, a characteristic gripping force on the order of ∼1 mN, and sufficient mucosal adhesion on the porcine tissue surface. In contrast, flipped bilayer actuators and single–layer PNIPAM or BSA–PEGDA controls showed positional drift and/or weak tethering under the same conditions (Figures S15–S17), supporting that robust retention arises from the integrated bilayer mechanism rather than either material alone. Collectively, these *ex vivo* results demonstrate strong, geometry–tunable gripping and retention under physiologically relevant motion and flow directions, supporting the feasibility of this platform for GI applications.

### *Ex vivo* retention probability under intestinal flow and surface morphology

To quantify the robustness of intestinal gripping beyond single demonstrations, we measured retention probability (%) under dynamic loading while accounting for tissue surface heterogeneity. Using the 6–arm edged snowflake as a representative geometry, nine bilayer grippers were placed at different locations along the porcine small–intestine surface to probe the effect of local morphology. We compared two bilayer thicknesses (0.5 mm and 0.25 mm per layer) and included two controls: reduced–size grippers (3 mm diameter) and single–layer BSA–PEGDA samples (Figure 6; Figures S18–S21).

**Figure 6.**
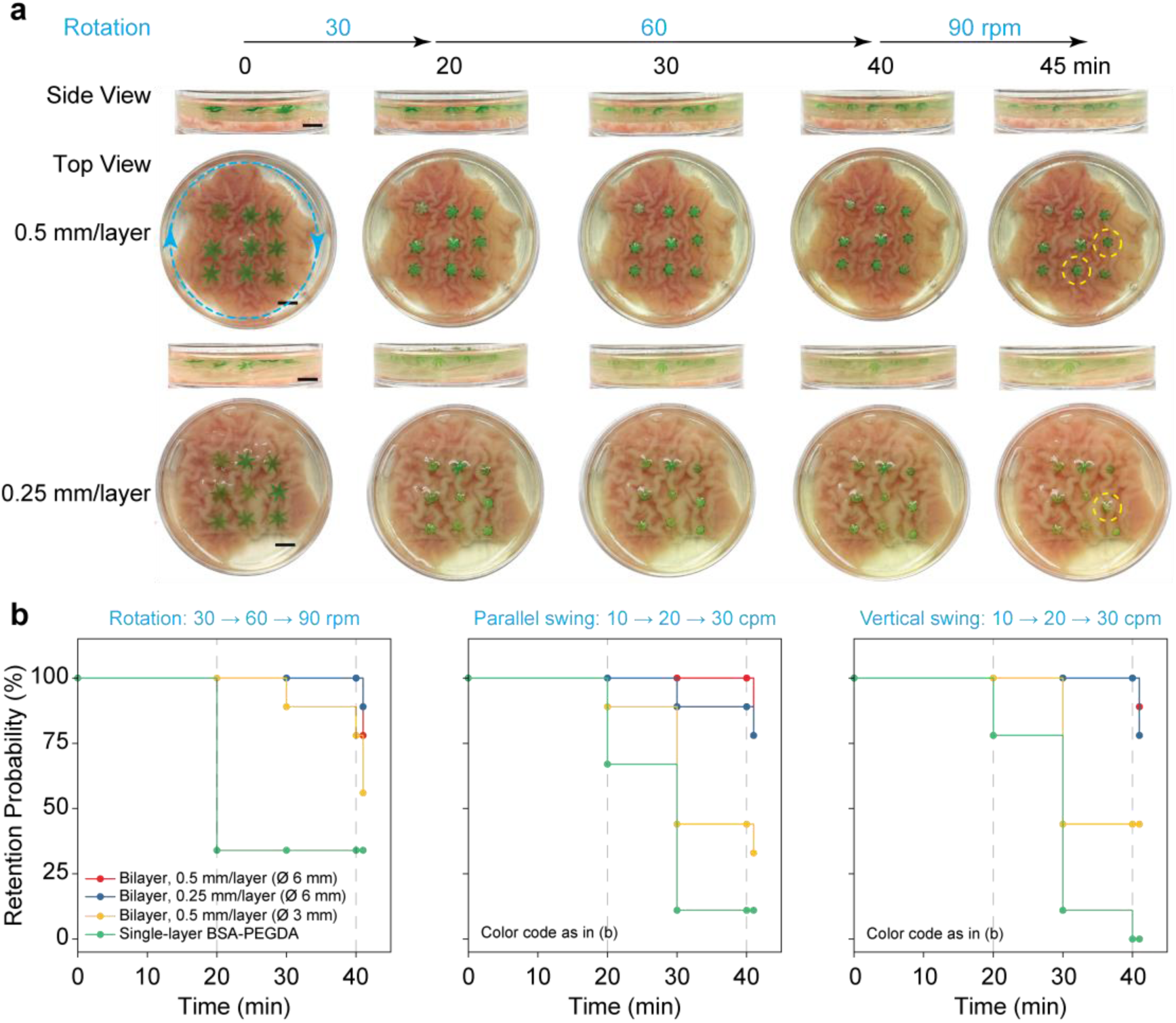
*Ex vivo* retention probability of PNIPAM/BSA–PEGDA bilayer grippers on porcine small intestine. **(a)** Representative time-lapse images of nine 6-arm edged snowflake grippers (Ø 6 mm) with two thicknesses (0.25 or 0.5 mm per layer) during rotation-mode testing; weakly tethered grippers are highlighted (yellow circles). **(b)** Stepwise retention probability profiles (n = 9 grippers per condition) under rotation (30 → 60 rpm; 90 rpm challenge) and swing (10 → 20 cpm; 30 cpm challenge) in directions parallel or perpendicular to plicae circulares. Final retention probabilities: rotation–78% (0.5 mm) and 89% (0.25 mm); parallel swing–78% (0.5 mm) and 78% (0.25 mm); vertical swing–89% (0.5 mm) and 78% (0.25 mm). Scale bar: 6 mm.

Across motion modes, bilayer grippers consistently outperformed controls (Figure 6). In the rotation mode, 0.5 mm bilayers achieved 78% retention (2/9 weakly tethered), whereas 0.25 mm bilayers achieved 89% retention (1/9 weakly tethered) (Figure 6). Although thinner grippers showed slightly higher retention, they were more prone to tilting due to increased flexibility, which can promote overbending and potential detachment during prolonged operation (Figure 6a). Controls performed substantially worse: the reduced–size group retained 56%, consistent with decreased contact area and incomplete anchoring, while single–layer BSA–PEGDA retained 33%, largely reflecting occasional mechanical trapping within plicae circulares rather than stable gripping (Figure 6b; Figures S20–S21).

Under plicae circulares–parallel swing, both bilayer thicknesses retained 78% (Figure 6b; Figure S18). One 0.25 mm gripper exhibited a marked positional shift at 40 min (highlighted in yellow), indicating reduced stability in this configuration. Controls again showed low retention (reduced–size: 33%; single–layer: 11%) (Figure 6b; Figures S20–S21). Under plicae circulares–vertical swing, 0.5 mm bilayers retained 89% and 0.25 mm bilayers retained 78% (Figure 6b; Figure S19), whereas controls remained poor (reduced–size: 44%; single–layer: 0%) (Figure 6b; Figures S20–S21).

Overall, these retention statistics demonstrate that standard–sized PNIPAM/BSA–PEGDA bilayer grippers (6 mm diameter) maintain robust attachment across heterogeneous intestinal surface regions and multiple flow directions, with performance strongly dependent on integrated bilayer function and adequate contact area. These results support the reliability of the platform for operation in dynamic GI–like environments and motivate future *in vivo* evaluation.

## Discussion

Stimuli–responsive hydrogels are versatile materials for biomedical technologies, spanning tissue engineering, drug delivery, soft robotics, and sensing [71]. In intelligent soft robotic systems, such hydrogels can integrate sensing, actuation, and functional output in a single, compliant platform [72]. Accordingly, thermoresponsive thera–grippers have been developed for minimally invasive drug delivery and tissue/cell sampling [33–35,37,69,73]. Enzymes have also been leveraged as endogenous cues to modulate hydrogel mechanics and morphology, enabling autonomous shape transformation and sustained functionality [30,74,75]. Multi–ink 4D–printing approaches further highlight how combining orthogonal stimuli (temperature, pH, enzymes) can increase adaptability in complex physiological settings [76].

Despite this progress, major barriers remain for translation of hydrogel thera–grippers: fabrication complexity and cost, grip performance and–most critically–the ability to achieve self–regulated actuation *in vivo*. Many systems rely on multilayer microfabrication with metallic components and multi–step alignment processes that limit biocompatibility and scalability [35,73,77]. Moreover, while multi–responsive actuators are common, shape recovery often requires externally staged triggers rather than intrinsic feedback. For example, a hinge–based gripper used sequential degradation of two spatially ordered biopolymer layers to close and then reopen, but depended on the external application of different enzymes and increased material complexity [74]. Likewise, enzyme–mediated cleavage strategies that impart degradability (e.g., disulfide cleavage leading to full degradation over hours) demonstrate valuable self–regulation concepts but typically require sophisticated synthesis and photolithographic fabrication [69]. Such external trigger dependence and fabrication burden constrain truly autonomous, task–adaptive behavior in dynamic physiological environments [78]. Grip force is a critical determinant for achieving long–term retention of devices within the GI tract. Recently, a bio–derived gripper fabricated from a euthanized spider and actuated by externally applied air pressure demonstrated a maximum grip force of ∼0.35 mN [79]. This result highlights both the inspiration drawn from various biological architectures and the potential to further enhance gripping performance through material optimization, particularly by leveraging swelling pressure mismatch and endogenous biophysiochemical stimuli.

Here, we introduce a bilayer hydrogel actuator that couples thermal actuation with enzyme–programmed mechanical feedback to enable GI gripping, localized trypsin delivery, and autonomous self–recovery. PNIPAM provides the thermoresponsive actuation layer, where the weight–based RSR governs the magnitude of thermally driven deformation and thus gripping capacity at 37°C (Figures 2a–c). The bilayer curvature is further tuned by the stiffness of the enzyme–programmed BSA–PEGDA layer: increasing BSA–PEGDA concentration reduces bending angle by mechanically constraining PNIPAM–driven deformation (Figure 2e). Encapsulating trypsin within BSA–PEGDA leverages intrinsic substrate–enzyme interactions to generate progressive self–softening of the protein–polymer network, evidenced by time–dependent decreases in Young’s modulus (Figure 3c; Figure S4a-c) and consistent with enzymatic regulation of protein–containing hydrogels [80,81].

Trypsin release measurements using FITC labeling show that matrix architecture governs release kinetics via coupled diffusion and network relaxation/erosion, supported by KP model fits (n > 0.45) under simulated intestinal conditions (Figure 3d,e; Figure S5) [55]. The release behavior shifts toward diffusion–dominated transport when enzymatic activity is suppressed (4°C) (Figures S6–S7), confirming the contribution of proteolysis–enabled matrix relaxation. Moreover, bilayer actuation accelerates release in denser BSA–PEGDA formulations, consistent with mechanical deformation promoting transport as digestion progressively weakens the network (Figure 3f,g; Figure S8) [82,83]. Functional delivery was validated by SDS–PAGE: released trypsin retained proteolytic activity and digested key whey proteins (β–lactoglobulin and IgG), confirming enzyme stability and functional release from the BSA–PEGDA matrix (Figure 3h,i; Figure S9) [56–61,84].

A key advance is the realization of autonomous actuation termination and shape recovery driven by the evolving mechanics of the enzyme–programmed layer. Bilayer grippers gradually reopened as trypsin–mediated digestion reduced the ability of the BSA–PEGDA layer to sustain interfacial constraint and curvature (Figure 4). Recovery kinetics depended on BSA–PEGDA formulation and protease availability: weaker networks opened earlier, while stiffer networks required prolonged incubation and/or exogenous protease to achieve measurable recovery (Figure 4a; Figures S10–S11). Importantly, self–release was demonstrated in a device–relevant gripper geometry, with opening accelerated in the presence of low external trypsin concentrations consistent with reported insufficiency regimes (Figure 4b; Figure S12) [63]. Together, these results establish a self–regulated workflow–grip, deliver, and autonomously disengage–without staged external triggers.

To evaluate performance under tissue–relevant conditions, we used fresh porcine small intestine as an *ex vivo* model [68]. Bilayer grippers generated characteristic gripping forces on the order of ∼1 mN, possessed excellent compatibility with small intestinal–relevant cell lines, and exhibited robust attachment under multiple motion modes and flow directions, whereas flipped or single–layer controls showed markedly reduced retention (Figures 5–6; Figures S13–S21) [69,70]. The compatibility of these devices is expected to further improve *in vivo* owing to the presence of a mucus layer that protects the epithelium [64,65]. This combination of mechanically effective gripping, superior compatibility, and programmed self–release addresses a central limitation of one–way GI–retentive devices–namely the risk of prolonged tissue intervention and abrasion–by providing a built–in “off–switch” after the therapeutic task is completed [43].

In a broader context, hydrogel soft robotics still faces challenges in material cost, self–regulated control, advanced fabrication, and biocompatibility of hybrid systems [62,72,78,85]. The PNIPAM/BSA–PEGDA(Trypsin) platform addresses these constraints by integrating (i) low–cost building blocks (NIPAM, BSA, PEGDA), (ii) endogenous logic (temperature and protease–mediated remodeling), and (iii) scalable, design–flexible DLP printing [86]. Compared with reported thera–grippers–including thermoresponsive polymeric/metallic systems, thermomagnetic grippers, and ionic shape–morphing microrobots–this approach emphasizes simplified fabrication, biocompatibility, and intrinsic functional autonomy [33,35,69,87].

Several limitations and opportunities remain. First, actuation speed is important for rapid anchoring *in vivo*; while grippers respond within ∼5 min (Video S1), further acceleration would improve practical utility. Second, the current design is intentionally single–use because protease–driven relaxation is irreversible; incorporating reversible physical crosslinks could enable multi–cycle actuation and multi–stage delivery [88,89]. Third, retention probability depends on local intestinal morphology and deployment location (Figure 6), motivating strategies for improved placement or higher–dose deployment to increase success probability *in vivo*. Finally, while DLP enables precise, customizable geometries and extends naturally to 4D printing concepts, material/ink constraints and bioink reusability remain practical challenges, particularly for UV/temperature–sensitive proteins and enzymes [86]. Maintaining protein function while ensuring print fidelity, and understanding how ink aging affects performance, will be critical for scalable translation.

In summary, the PNIPAM/BSA–PEGDA(Trypsin) bilayer actuator provides a proof–of–concept strategy that couples thermal actuation with enzyme–programmed mechanical feedback to achieve GI gripping, localized enzymatic supplementation, and autonomous self–release–bridging conventional oral administration and more intelligent, self–regulated therapeutic systems for GI applications.

